# Population Dynamic Regulators In An Empirical Predator-Prey System

**DOI:** 10.1101/2020.08.23.263376

**Authors:** Anna S. Frank, Sam Subbey, Melanie Kobras, Harald Gjøsæter

## Abstract

This paper investigates stability conditions of an empirical predator-prey system using a model that includes a single delay term, *τ*, in description of the predator dynamics. We derive theoretical conditions on *τ*, in terms of other model parameters, and determine how changes in these conditions define different stability regimes of the system. We derive optimal model parameters by fitting model to empirical data, using unconstrained optimization. The optimization results are combined with those from the theoretical analysis, to make inference about the empirical system stability.

Our results show that Hopf bifurcation occurs in the predatory-prey system when *τ* exceeds a theoretically derived value *τ*\* > 0. This value represents the critical time for prey availability in advance of the optimal predator growth period. Set into an ecological context, our findings provide mathematical evidence for validity of the match-mismatch hypothesis, for this particular species.

## 1 Introduction

If *x* ∈ ℝ_≥0_ and *y* ∈ ℝ_≥0_ are state variables that represent population indices (e.g., abundance) respectively, of prey and predator, then the classical Lotka-Volterra Model (LVM) [1, 2] description of the system, 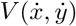, is defined by

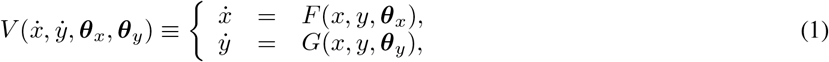

where *F*: ℝ_+_ → ℝ_+_ and *G*: ℝ_+_ → ℝ_+_ are continuous functions, and ***θ**_x_* ∈ ℝ^*n*^ and ***θ***_*y*_ ∈ ℝ^*m*^ are sets of parameters associated with *x* and *y*, respectively. Several theoretical analyses in the current literature (see e.g., [3, 4, 5]) determine how ***θ**_x_* and ***θ***_*y*_ influence system dynamics.

Following e.g., [6], we can define

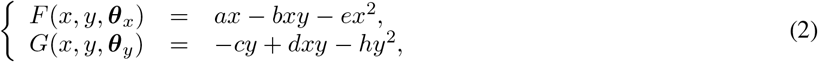

then ***θ**_x_* ≡ {*a, b, e*}, and ***θ***_*y*_ ≡ {c, d, h}. In (2), the term *ax - ex*^2^ defines a logistic growth with a limiting carrying capacity *a/e* (see e.g., [7]), and *bxy* is loss in prey biomass due to predation, also known as the functional response [8]. The predator dynamics is determined by a natural death rate term *cy*, population decrease due to intraspecific competition *hy*^2^, and *dxy*, which defines the biomass gain through predation.

Since the parameters in ***θ***_*x*_ and ***θ***_*y*_ determine trajectory of the state parameters, their definition is reflective of the underlying ecological assumptions. Thus, one may constrain any *θ* ∈ **θ**_*x*_ (similarly for ***θ***_*y*_) to a scalar domain in ℝ or to the domain of some continuous function in ℝ.

Seasonality is an important characteristic feature of e.g., boreal and arctic environments for population growth [9, 10] and an obvious driver of the system (see e.g., [11]). For example, the authors in [12] assumed that seasonality has continuous time effect by defining growth rates, both *a* and *c*, as smooth sinus functions of time. However incorporating such information in the population dynamics (of either predator and/or prey) may be generally, non-trivial, especially as different functional expressions of the seasonality may lead to different scenarios of the population trajectory [13].

Another ecological consideration is the fact that a time delay exists between when prey is ingested, until it is converted to predator biomass. The simplest approach to addressing this consideration is to introduce a constant delay *τ* > 0 in the functional response term of the predator [14], so that *G*(*x, y, **θ**_y_*) is redefined as

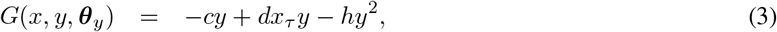

where in general, we use the notation *x_τ_* ≡ *x*(*t* − τ).

Though a time-lag to account for gestation may be ecologically sound, the literature shows that time-lags have a destabilizing effect on the system dynamics (see [15]). Thus, on one hand, a system with a stable equilibrium may transition to an unstable and/or oscillatory population as the lag increases, and may even cause a bifurcation into periodic solutions. On the other hand, for the predator, the functional response term in (2) has a stabilizing effects on the population dynamics. For an empirical population where the integrated system may form the basis for inference about the population dynamics, an accurate estimate of *τ* is therefore important. Deriving the value of *τ* from empirical observations is, however, non-trivial. Predator-prey systems do not act in isolation of their environment. The environmental factors (biotic and abiotic) may act on different, individual time-scale resolutions, and feedback loops (i.e., different delay terms) to dictate the population dynamics (see e.g., [16]). Hence the system of equations with constant delay *τ* may be an approximation of one with infinitely distributed delays.

When data on both predator and prey are available, the parameters associated with *F* and *G* in (1) may be determined (as for example, in [17]) by numerical optimization. If 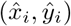, *i* = 1,…, *n*, represent a set of empirical observations over *n* discrete time steps, and 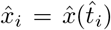 (similarly for 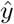), one may define the initial conditions for a system by setting 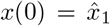, and *y*(0) = *y*_1_. One challenge is that, for the DDE system, *τ* and *x* at *t* ∈[-*τ*, 0) must be known. Deriving these values from empirical observations may be non-trivial. Given fixed observation time intervals 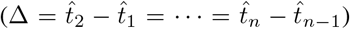, [0, Δ) and [Δ ∞) are two feasible intervals for *τ*. If the problem is formulated (using e.g., a constrained optimization approach) to include the estimation of *τ*, the derived solution will depend on the chosen feasible interval. In practice, empirical data are however not always available to test validity of ecological assumptions. In absence of data however, it would be especially restrictive to constrain parameter domains of the model. Also with partially available empirical data (only for either predator or prey, but not both), the estimation problem in (1) is challenging. Then auxiliary information is needed to validate the derived trajectory of the missing component.

This paper investigates the dynamics of a predator-prey system, where empirical data on the predator, but not the prey is available. Auxiliary information for validation of prey dynamics are measurements of the annual average weight of the predator. The predator is the Barents Sea (BS) capelin, which is a short-lived (1–4 years) fish species that spawn only once in their lifetime and die [18]. Capelin preys on the lowest level species (zooplankton species), which usually advected into the BS through Atlantic and Arctic water influx [19, 20]. Its selected prey size and type is age-dependent [21]. Capelin is itself, prey to other species at higher levels in the food chain, and preferred prey for the Northeast Arctic cod [22]. Several episodes of extreme capelin biomass decline (collapse) have been observed during the last three decades. Large predation (top-down effect) from other species (during crucial capelin life stages) [23, 24] has served as one explanation for these episodic events of collapse. Another explanation that has been reported in the literature is that food availability (bottom-up effect) regulates capelin population dynamics (see e.g.,[21]). Although the latter effect is considered to be of less relevance than the top-down effect, statistical modeling and analysis showed that a bottom-up regulation of capelin biomass dynamics is significant [25].

The goal of this paper is to investigate whether the capelin biomass dynamics may be reconstructed (including episodic events of extreme decline) by solely considering a bottom-up regulation process. The paper adopts the model definitions in (2) and (3), and uses unconstrained optimization to estimate the parameters of the system. We make inference about the system dynamics using the derived parameters. The manuscript is organized in the following way. Section 2 gives a summary of the observation data on which the modeling is based. Section 3 revisits the mathematical models, and presents their theoretic dynamical system analyses. This section also presents a formal definition of the optimization problem, whose solution yields the system parameters. The theoretical results inform inference on the predator-prey behavior, on the basis of the derived parameters. Section 4 gives an overview of the numerical experiments, whose results are presented in Section 5, and discussed in Section 6. Our conclusions and discussion about possible limitations of the results, are presented in Section 6.1.

## 2 Data sources

From annual scientific cruises in the BS during September, data on species abundance, spatial distribution and demography have been obtained, since 1972 [26]. The species abundance indices (length, weight, age, numbers) are converted into age-specific biomass.

Figure 1 shows the biomass of capelin from 1972 to 2013 (for the indicated ages), with notable stock collapses in 1985–1989, 1993–1997, and 2003–2006 [22]. This paper uses time series data of the age-structures capelin biomass (Figure 1) as predator data.

**Figure 1:**
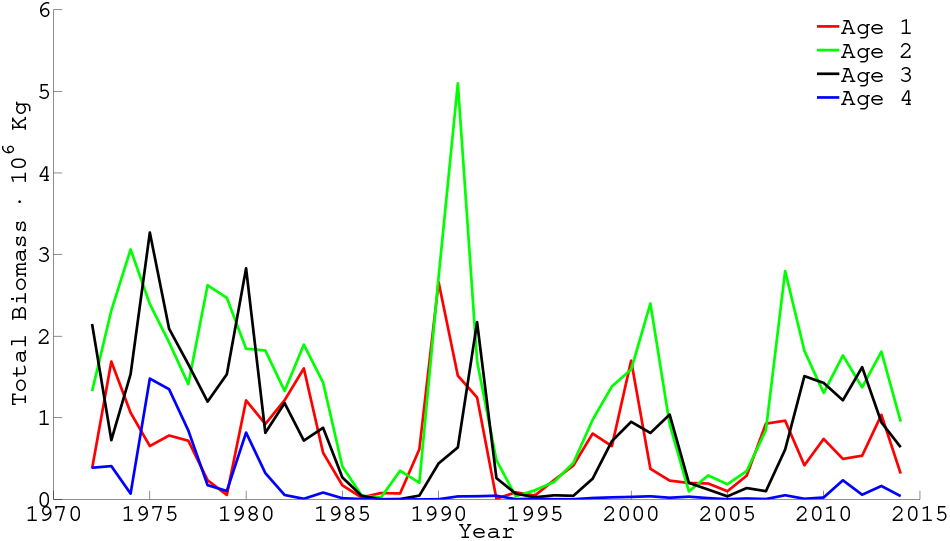
The capelin biomass data

For parameter estimation and modeling of predator dynamics, we use a time-series of capelin biomass data from 1990-2018. The data for age-4 capelin is excluded from the modeling because the observations are highly uncertain, infrequent, and low. Capelin biomass data are not reliable before 1983 [18], and capelin biomass was extremely low between 1986-1989. Hence our simulations (and parameter estimation) start in 1990, when the stock has recovered [27].

We have used the term *prey* to collectively describe all capelin prey, with a wide repertoire from Arctic copepod species to Atlantic krill [28]. This decision is influenced by the following considerations. Firstly, we have considered the prey data as unavailable partly because we are unable to pin down the exact prey type. Secondly, the intensive feeding season for capelin is July–October [26]. However, the scientific survey that collects data on the abundance of main capelin preys occurs at the end of this season. We consider this data therefore, as reflective of the residual prey abundance. The average weight of capelin has thus been used as a proxy for the validation of of the modeled prey trajectory, 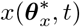.

All data used in this manuscript have been obtained from the database of the ICES Working Group on the Integrated Assessments of the Barents Sea [29].

## 3 Model and Theoretical Analyses

The main mathematical model description and theoretical analyses are presented in this section.

### 3.1 The general predator-prey model

We use the system in (1), where for *τ* ≥ 0, we define

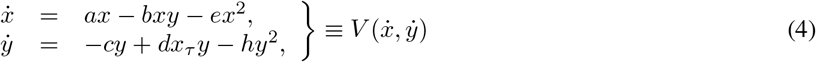

which are adaptations from [6] and [14].

### 3.2 Theoretical analyses of system dynamics

#### 3.2.1 Coexistence and conditions

We use (4) to derive the non-trivial equilibrium point, *P*_*_ ≡ (*x*_*_, *y*_*_), of the system 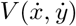, which is given by

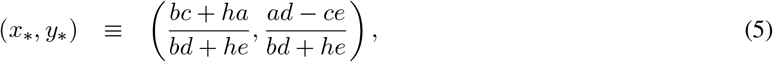

where

- Condition 1 (**C1**): *ad* > *ce*

We introduce the transformations

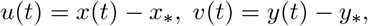

to derive (6) from (4).

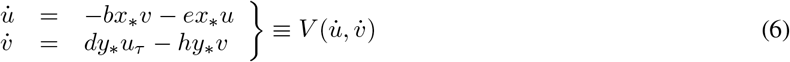

Equation (7) gives the characteristic equation for 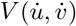,

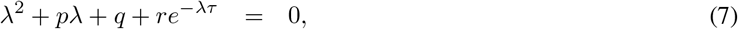

where *p* = *ex*_*_ + *hy*_*_ ≥ 0, *q* = *hex*_*_*y*_*_ ≥ 0, and *r* = *bdx*_*_*y*_*_ ≥ 0. The equilibrium *P*_*_ is stable if all roots of (7) have negative real parts. For *τ* = 0, we derive the characteristic equation (8), with discriminant *D* = *p*^2^ − 4(*q* + *r*).

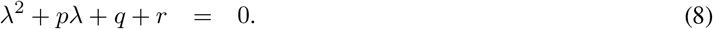

Then the roots of (8) have (i) always negative real parts when {*p* > 0 ∧ *D* < 0}, (ii) always positive real parts when {*p* < 0 ∧ *D* < 0}, and (iii) at least one positive root for {*p* < 0 ∧ *D* ≥ 0}. The constraining conditions are defined as

- Condition 2 (**C2**): (*ex*_*_ + *hy*_*_) > 0 ∧ (*ex*_*_ − *hy*_*_) < 4*dbx*_*_*y*_*_, and
- Condition 3 (**C3**): (*ex*_*_ + *hy*_*_) < 0.

#### 3.2.2 Stability and bifurcation analysis

If we define *λ* = *iw* as a root of (7), we derive (9)–(10),

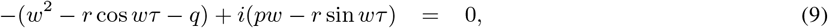

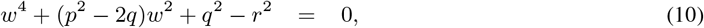

which come from separating the real and imaginary parts. We introduce

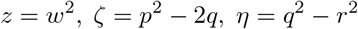

into (10), to arrive at (11)

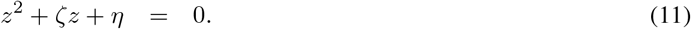

Note that *ζ* = *p*^2^ − 2*q* > 0 since (*p*^2^ − 4*q*) = (*ex*_*_ − *hy*_*_)^2^ ≥ 0, and the discriminant of (11), *D* > ζ^2^, for *q* < *r*. Hence, when *q* < *r*, has a positive root, *z*_0_, for *z* ∈ (0, *∞*), and no positive root when *r* ≤ *q*. The positive root of (10), *w*_0_ is then given by 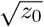, i.e.,

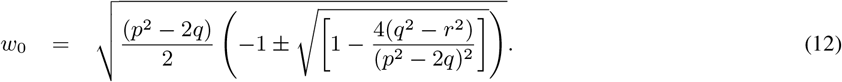

Using the real part of (9), we define for *κ* = 0, 1, 2,…

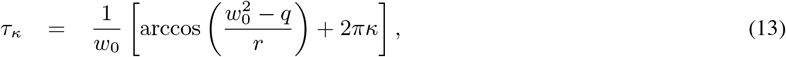

and note that (7) with *τ* = *τ_κ_* has a pair of purely imaginary roots, ±*iw*_0_ for every *τ_κ_*. We investigate the transversality condition, expressed as Lemma 1.

##### Lemma 1

*Let λ*(*τ*) = *α*(*τ*) ± *iw*(*τ*) *be the root of (7) near τ* = *τ_κ_, such that α*(*τ_κ_*) = 0, *and w*(*τ_κ_*) = *w*_0_ *for κ* = 0, 1, 2,…, *Then if q* < *r*,

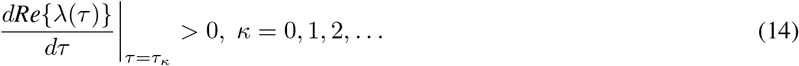

**Proof 1.***Rewrite (7) with explicit dependence of λ on τ, i.e., λ ≡ λ*(*τ*), *as*

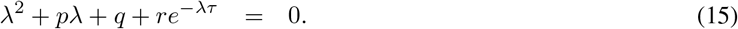

*Then*

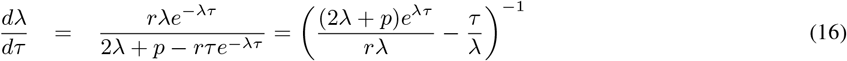

*However*,

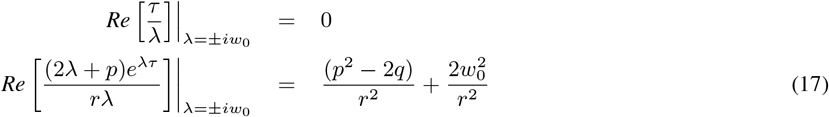

*since from (9)*,

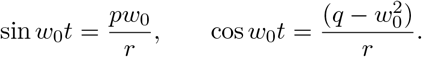

*Finally, substituting for w*_0_ *from (12), we deduce*

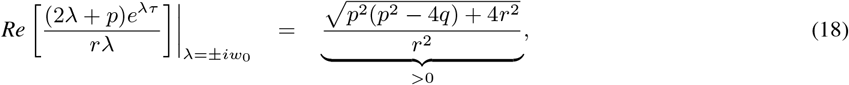

*which completes the proof*.

##### Theorem 1

*Let* **C1** *and* **C2** *prevail. Then all the roots (7) have negative real parts for τ ∈* [0, *τ*_0_), *and P*_*_ *is asymptotically stable for τ* ∈ [0, *τ*_0_). *The system defined by (2) undergoes Hopf bifurcation at P*_*_ *when τ* = *τ_κ_, κ* = 0, 1, 2,…

##### Theorem 2

*Let* **C1** *and* **C3** *prevail. Then (7) has at least a root with positive real part for τ* ∈ [0, *τ*_0_), *and P*_*_ *is unstable for τ* ∈ [0, *τ*_0_). *The system defined by (2) undergoes Hopf bifurcation at P*_*_ *when τ* = *τ_κ_, κ* = 0, 1, 2,…

### 3.3 The optimization problem

The assumption that discrete empirical observations exist only for the predator, and not the prey, defines the optimization problem. Our goal is to determine the system parameter sets that minimize the discrepancy between modeled predator biomass and empirical data.

Define 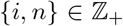 such that *i* = 1,… *n*, and let 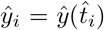 represent empirical observations of *y* over *n* discrete time steps 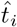. Note that *y* and *x* are coupled (through the functional response). Furthermore, since we assume no data exists on 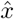, the initial condition *x*(0) and delay term *τ* must be estimated as part of the optimization procedure. The trajectory of *y* will also depend on *x*(0) and *τ*. Hence we write *y*(***θ**_x_, **θ**_y_, x*(0), *τ, t*), and define Problem 3 as the general optimization problem.

#### Problem 3

*(The optimization problem)*

*Determine **θ*** ≡ {***θ***_*x*_, ***θ**_y_, x*(0), *τ*}:

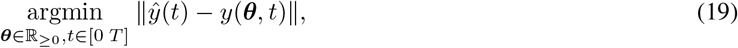

*where* 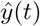 *is known at* 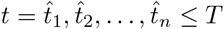.

## 4 Description of numerical simulations

Determining the trajectories of the predator-prey system combines the problem of integrating the system of coupled DDEs, and deriving the solution, ***θ***\*, of the optimization Problem 3. We use a numerical approach for integrating the DDE system, as for most of such coupled systems, finding closed form solutions is non-trivial [30].

For the coupled DDE system, we use the Matlab dde23 algorithm, which is based on the ode23-solver, using the RKBS(2,3) method. Theoretical and computational details about dde23 can be found in [31]. We set 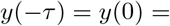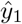, and *x*(−*τ*) = *x*(0) = *x*_0_. Since *τ* and *x*_0_ are unknown, they are included in the parameter set ***θ*** (see Problem 3).

We used the fminsearch algorithm in Matlab to derive the unconstrained model parameters. The fminsearch algorithm uses the Nelder-Mead algorithm [32] to compute the unconstrained minimum of a given objective function. For a predefined tolerance *ϵ*, we consider the algorithm to have converged after *J* iterations when the change in the objective function Δ*f*(***θ**_j_*) < *ϵ*, for *j* ≤ *J*. For each candidate solution, ***θ**_j_* (*j* = 1,…,*J*), spline-interpolated values of *y*(***θ**_j_*, *t*) at *n* discrete (observation) points 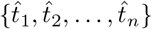 are obtained.

The objective function is then simply defined by (20).

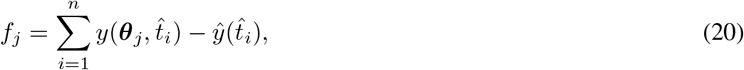

where, *f_j_* = *f* (***θ**_j_*). Based on the analysis presented in Section 3, we obtain system dynamics by analyzing the set of optimized parameters.

Finally, we validate the dynamics of the modeled prey trajectory, 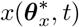, by comparing it to data on the average weight of capelin biomass.

## 5 Results

This section presents results from our numerical simulations for the DDE model. We adopt the following notations, some of which has been used in Section 3, but are repeated here for the sake of completeness and to ease readability:

- ***v*** = (*a, b, c, d, e, h, τ*)
- ***v**^o^* ≡ optimized parameter set
- *τ* * ≡ critical Hopf bifurcation parameter
- *x*_0_ ≡ Initial condition for prey
- Let *τ*^†^ define an arbitrary value such that:

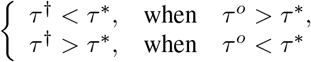

Table 1 shows results for our numerical simulations when *τ* > 0. We used the optimized estimate of *τ* and results in Theorem 1 to calculate *τ_κ_, κ* = 0, 1, 2,…, though only *τ*_0_ and *τ*_1_ are shown here. Observe that results from the optimization show that it is Theorem 1 (and not Theorem 2) that applies to our empirical setting. Although we are able to calculate *τ_κ_, κ* ≥ 0, only *τ*_0_ presents a biological realistic scenario, since capelin has a life-span of three years. The results presented in this section are therefore limited to the model dynamics for varying *τ* - values around *τ*_0_. Figure 2a.–b. shows the simulated prey and predatory biomass dynamics, as well as comparison between modeled and empirical predator biomass data. The DDE system is capable of replicating the empirical observations. Figure 2c. shows a consistent, temporal synchrony between the modeled prey biomass, and the average weight of age-3 capelin. Observe that, as expected, there is a slight time-lag between the scaled total prey biomass, 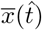, and the scaled predator weight, 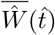. Coherence between modeled prey dynamics and average predator weight is less pronounced for ages 1 and 2 capelin, compared to the results for age-3 (see Figure A.1 in the appendix).

**Table 1:** Parameter estimation results using data from 1990–2018 (age-1 and 3), and 1989-2018 (age-2). The optimization algorithm failed to converge for age-2 when data from 1989 was excluded – see discussion under Section 6

| Age | $v^o = (a, b, c, d, e, h, \tau_0^o)$ | $x_0$ | $\tau_0^*$ | $\tau_0^\dagger$ | $\tau_1$ |
| --- | --- | --- | --- | --- | --- |
| 1 | (1.1747, 2.0315, 0.750, 0.300, 0.0380, 0.3747, 0.1268) | 16.2625 | 0.3204 | 0.5251 | 6.6966 |
| 2 | (0.4893, 0.3974, 0.9874, 0.2649, 0.0070, 0.1488, 0.0556) | 13.1306 | 0.3860 | 0.7163 | 9.1301 |
| 3 | (0.5398, 0.8562, 1.0310, 0.2325, 0.0095, 0.0594, 0.1070) | 8.9218 | 0.1473 | 0.1876 | 8.7989 |

**Figure 2:**
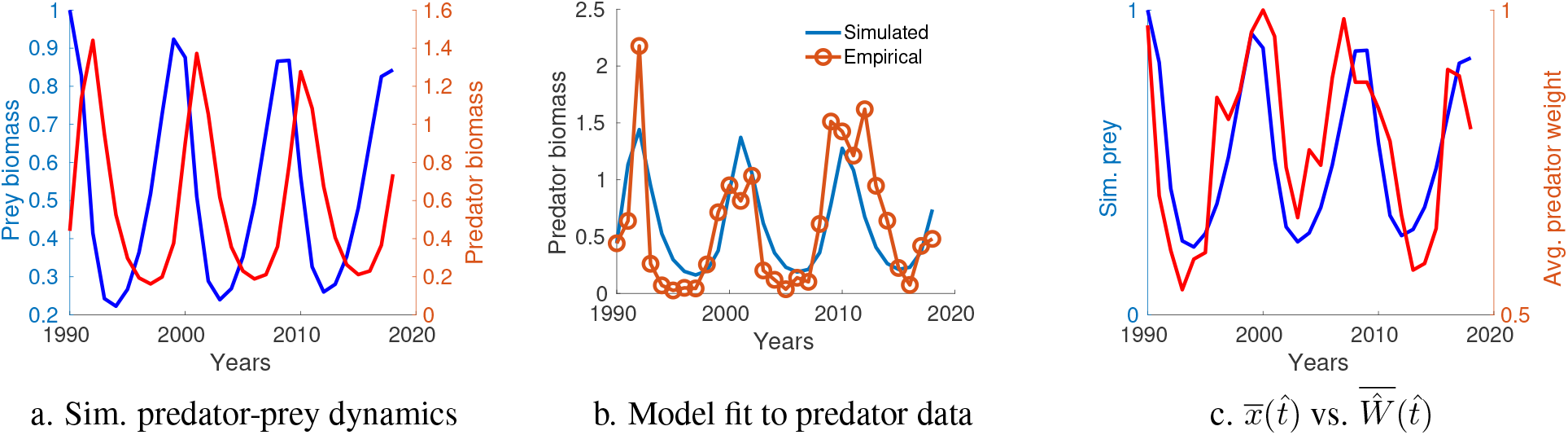
DDE optimization results – Simulated predator-prey dynamics (a.), Model fit of predator dynamics to empirical data for age-3 capelin (b.), Scaled total biomass of prey, 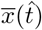, and scaled predator weight, 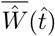, in the period 1990–2018 (c.)

The first column in Figure 3 shows fit of the DDE model to the data with the indicted *τ* values. The second column shows the corresponding phase-plane diagrams.

**Figure 3:**
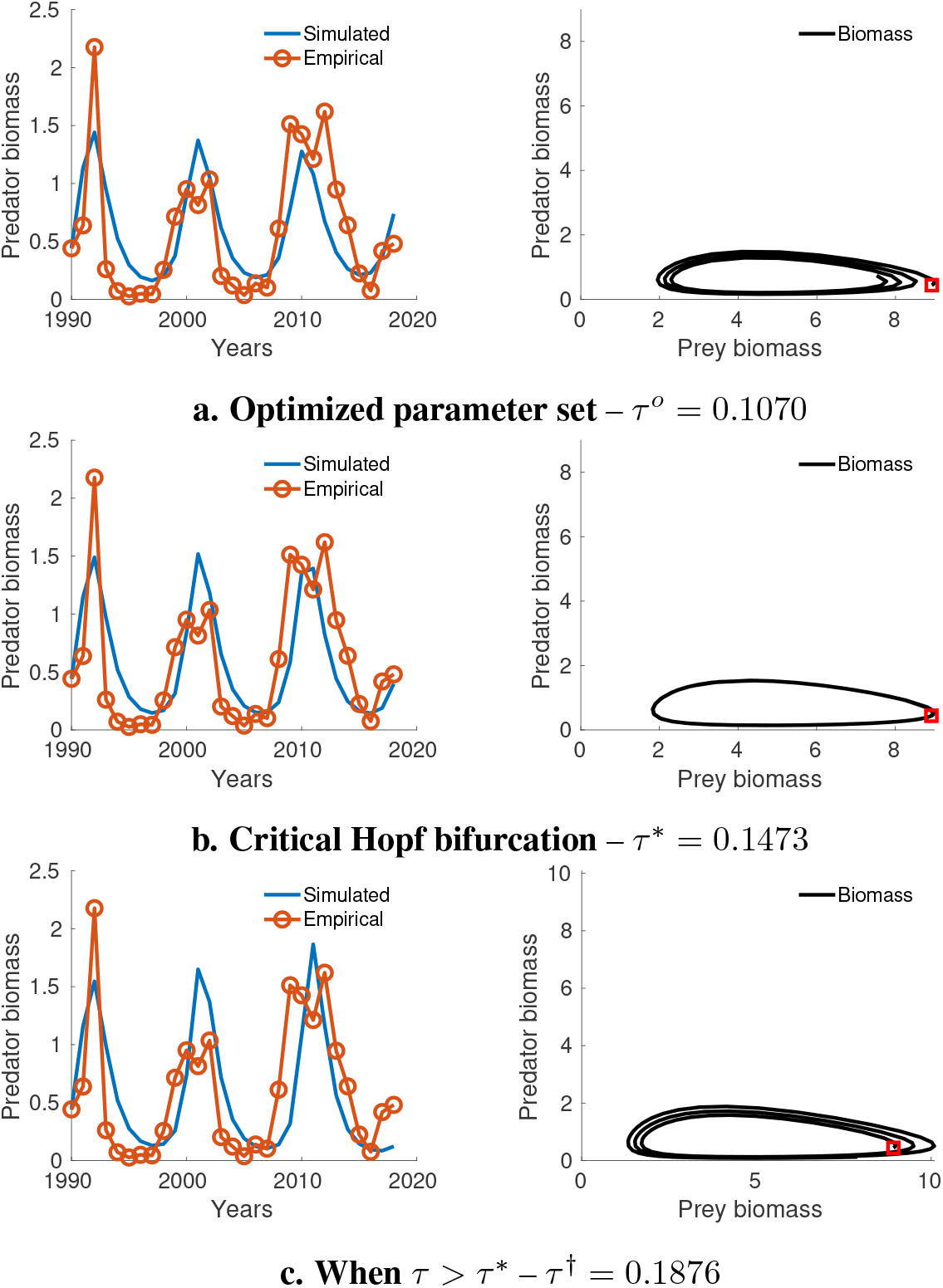
*τ*-bifurcation analysis – **Left column:** Model fit to data. **Right column:** Phase-plane plot of the predator-prey dynamics. Observe the phase-plane dynamic transitions from (a) Asymptotic stable (*τ^o^*) → (b) A limit cycle (*τ \**) → (c) Asymptotic unstable (*τ*^†^). The red square in the phase-plane plot is the origin of the time-series.

## 6 Discussion

This paper analyzed the dynamics of an empirical marine predator-prey system, both theoretically and numerically, based on Lokta-Volterra model formulations, where data on the prey is absent. We have used auxiliary information about the predator to validate the simulated prey biomass dynamics. Although, full simulation results exist also for age-1 and age-2 capelin (Table 1), graphical results and discussions were focused on age-3 capelin. This has been done, partially for the sake of brevity. But most significantly, because the results for age-3 capelin are coherent and easier to interpret. The inability to validate the modeled age-1 and age-2 prey dynamics with data on averaged predator biomass weight might indicate that the younger age-groups are not entirely regulated by bottom-up process, in contrast to age-3 capelin, and this warrants further investigation. However, a alternative, and perhaps, more plausible explanation may exist for our ability to better fit the age-3 capelin.

If we define

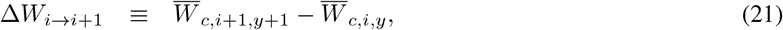

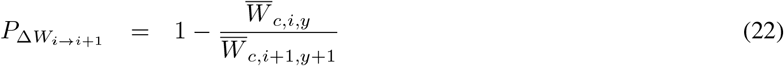

where 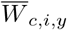 is the average weight of fish cohort *c* at age *i* in year *y* (and similarly for 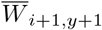, then for *i* = 1, 2, 3, we derive three groups of Δ*W*_*i*→ *i*+1_. Figure 4 shows a Boxplot of data for the three groups of Δ*W*_*i*→ *i*+1_, for cohorts from 1990-2008. This figure shows the highest variability in average weight from age 1 → 2 (approximately 68%), and lowest for 3 → 4 (approximately 15%). In general using average weight by age could be problematic for fish of age-2 and higher, as the component of the stock that is measured is affected by the combination of length-dependent maturity and total spawning mortality. The largest age-2 fish measured in autumn, will mature, spawn, and die, before next autumn cruise. Consequently, these do not form a part of the cohort of age-3 fish whose mean weight are measured the following autumn. This may result in underestimation of annual mean weight for fish older than two years which, consequently, weakens the dynamic link between average weight and prey abundance. The underestimation (and its consequence) will be expected to increase with age [33]. On the other hand, since weight by age represents accumulated growth throughout the year up to the time of sampling, the accumulation process can potentially lead to a smoothness of the data uncertainty. Consequently, the link between food and weight at age becomes stronger (as our results indicate) with increasing age.

The average weight from age-3 to age-4 capelin appears to be relatively stable, as the mean percentage change is approximately 16% (compared to 68% for age-1 to 2). Thus results from Figure 4 may explain why prey abundance and average age-group weight may be more consistent with increasing age-group, especially when using a model that does not discriminate between prey type. For younger age-groups, more stochastic (e.g., defined by stochastic differential equations) prey model(s) may be required. Consequently, our modeled prey dynamics may represent a single realization of several stochastic dynamic prey trajectories.

**Figure 4:**
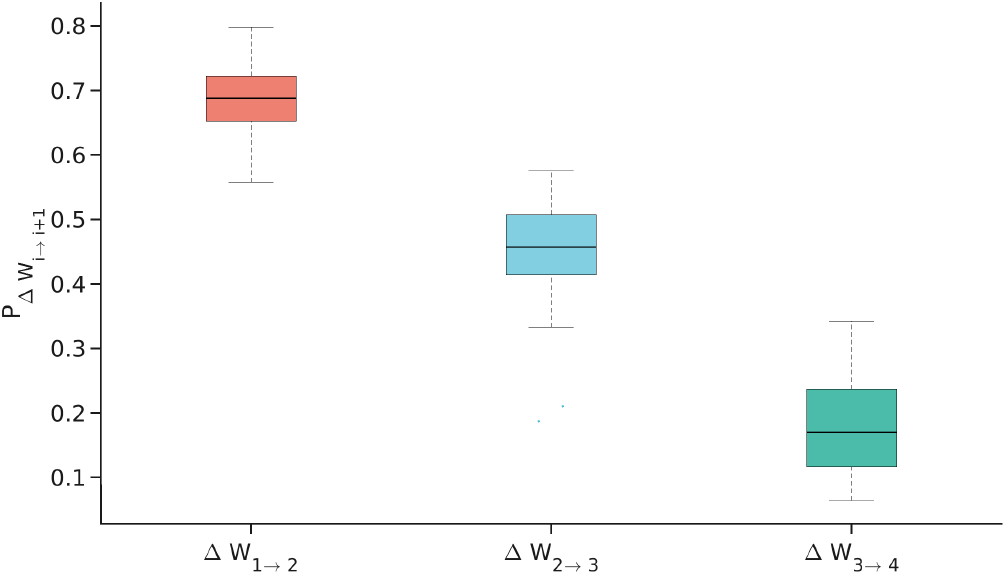
Boxplot of fractional change in average weight, *P*_Δ*W*_i→i+1__, of cohorts at age *i* (in year *y*) and age *i* + 1 (in year *y* + 1) for *i* = 1, 2, 3., using data from 1990–2018.

In general however, we see good model fit to empirical data, for stable and unstable system equilibria (see Figure 3).

Since **C1** and **C2** apply, Theorem 1 defines the condition for stability of the predator-prey system. This is confirmed by the results in Table 1 and Figure 3, where for *τ* ∈ [0, *τ*_0_], the system dynamics is stable, but becomes unstable when *τ* = *τ ^†^* > *τ*_0_.

Table 1 shows that the optimal delays (*τ*^0^) and critical values (*τ*\*), are age-class dependent. Put together, these delay values define a time-window, Δ*τ* ≡ [*τ*^0^, *τ*\*], for stability of the predator-prey system. From (2), we note that for *{c, d, h}* ≥ 0, the growth rate *G*(*x, y, **θ**_y_*) at any time *t*, satisfies (23),

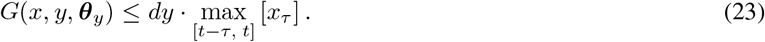

Hence, predator growth is optimized when *x_τ_* is maximum in [*t* − *τ*, *t*], for *τ* ∈ Δ*τ*, and that this time-window is age-dependent. For the optimal time-delay window (in weeks), we observe that Δ_1_*τ* ≡ [8, 16], Δ_2_*τ* ≡ [3, 16], Δ_3_*τ* ≡ [4, 8], where Δ_*j*_ is the time-window for predator of age *j*.

Combining (23) and the results in this manuscript translate, in ecological terms, to mean that the growth rate (and stability) of the predator at any time *t* are determined by the size of prey biomass within *t* - Δ_*j*_ (for *j* = 1, 2, 3), weeks in advance. These differences in Δ*τ* is reflective of the age- or length-specific feeding needs and prey types for capelin (see discussion in [21]). For instance, capelin forages mainly during the autumn, and lipid reserves built during this time, sustain juvenile capelin during winter, when it does not feed [26].

We extend interpretation of our results in an ecological context by firstly noting that feeding among fish is a dynamic process. A newly hatched capelin larva would, for instance, depend on availability of food objects of a narrow size spectrum. If such food (for instance eggs and young stages of crustacean plankton) is not available soon after the capelin larvae are hatched, they will have minimal chance to survive. In theory similar mechanisms may exist also for older fish, but since they can choose among a wide range of food objects, they are much less prone to lack of suitable food. Adult fish is also extremely flexible as to when food is available; they may survive for long periods without eating anything, thanks to their low resting metabolism.

Our results show that the dynamic optimal growth rate (not necessarily the maximum) at any time *t*, is determined by the size of prey abundance in the time-window *t* - Δ_*j*_ in advance. Note that what constitutes maximum predator growth rate is not only dependent on the prey abundance *per se*, as growth rates may be dictated by other factors, e.g., ratio of prey to predator biomass, handling time, and optimal spatial overlap between predator and prey.

Supposing Δ*T* ≡ [*T*_0_, *T_f_*] represents an arbitrary time interval during a feeding season where prey is distributed over a spatial region **Ω** ∈ ℝ^2^. Let Ω(*t*′) ∈ **Ω** be a spatial patch with highest prey density (e.g., plankton bloom) at time *t^I^*. Then optimal growth conditions observed at time *t* ∈ Δ*T*, is a consequence of predator-prey overlap on patch Ω(*t*′), at *t*′ = *t* - Δ_*j*_. An ecological significance of our time-windows deals with the match/mismatch hypothesis (MMH) [34], which asserts that dynamical variations in a population are driven by the relationship between its phenology (i.e., timing of population seasonal activities) and that of the immediate lower trophic level species. Normally, the MMH is limited to deal with mortality of fish larvae during their critical early life stages. However, the concept may be broadened to include other life history parameters than mortality, e.g. growth and maturation, and the original temporal aspects may be broadened to include also spatial aspects of match and mismatch, as exemplified above.

Though the MMH hypothesis has gained acceptance, the literature lacks empirical evidence for its validity (see discussion in [35]). Our derived time windows, Δ_*j*_, provides a possible link between capelin phenological growth and that of its prey. To the authors’ knowledge, this is the first time such evidence of the MMH has been provided.

Although our results show that the observed cyclic variation in biomass of age-1 and older capelin is consistent with the hypothesized bottom-up regulation, this does not preclude the existence of top-down regulation both on early life stages (reviewed by [36], and not dealt with in our modeling) or among adult capelin. The importance of bottom-up and top-down regulation might shift among various life history stages of capelin.

### 6.1 Conclusions

Our modeling results shed light onto the regulatory effect of prey on BS capelin biomass dynamics. Results from theoretical analyses and numerical simulations were consistent. We could also show that the simulated prey dynamics are perfectly synchronous with annual average predator weight, and have thus validated our model formulation of the prey.

Three key ecological highlights result from our analyses. Firstly, we have presented results in support of a bottom-up regulation for the biomass dynamics of capelin age-1 and older. Secondly, based on combination of theory and simulations, we have identified time-frames for predator-prey overlap, which lead to optimal predator growth and stability of the predator-prey system. The identified time-frames differ for different age groups, and probably reflect age-specific feeding habits of the predator. Thirdly, we have provided evidence, perhaps for the first time, of MMH applicability to capelin and its prey.

Our results present evidence to show that prey have strong regulatory effect on the biomass dynamics of predators of age-1 and older. However, we cannot infer from our results whether this can also partly explain the episodic collapses seen in the capelin biomass dynamics. In our opinion, such inference, should be based on findings from this paper, combined with analyses of other biophysical information in space and time. However, this undertaking is beyond the scope of this paper and will therefore be investigated in a sequel paper.

## Acknowledgements

SS is grateful to the Fulbright Foundation (Norway) for funding his research at Cornell University, where work on this manuscript was initiated.

## A Appendix

**Figure A.1:**
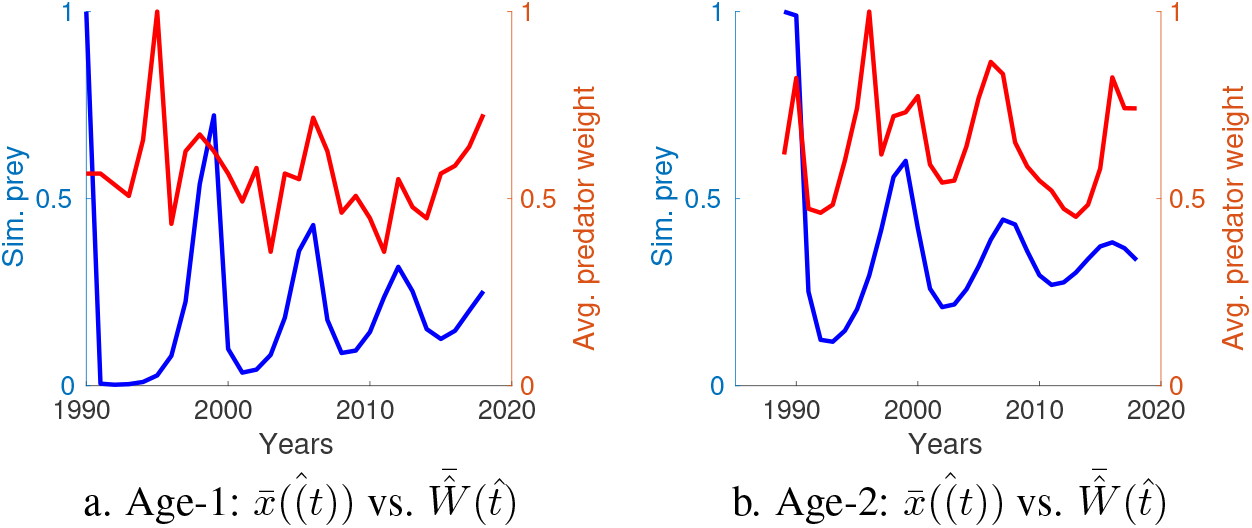
Scaled total biomass prey, 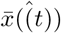 and scaled predator weight, 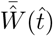, for predator of age-1 (1990–2018) (a.), and age-2 (1989–2018) (b.)

